# Progranulin deficiency aggravates aging-induced vascular injury

**DOI:** 10.1101/2025.11.25.690611

**Authors:** Renata de Azevedo Melo Luvizotto, Andre F Nascimento, Gustavo F Pimenta, Rafael M Costa, Shubhnita Singh, Juliano V Alves, Wanessa Awata, Angela Russ, Adeyeye Haastrup, Rolando Cuevas, Parya Behzadi, Cynthia St Hilaire, Justin Roberts, Ariane Bruder, Thiago Bruder-Nascimento

## Abstract

**Significance:** Vascular aging is a major contributor to cardiovascular disease, yet the molecular mechanisms of age-associated vascular dysfunction remain incompletely defined. This study reveals a critical role for progranulin (PGRN) in regulating vascular senescence, function, and remodeling during aging.

**Methods:** We assessed PGRN expression in human and mouse arteries and senescent vascular smooth muscle cells (VSMCs). Functional vascular studies were performed in PGRN-deficient (PGRN⁻/⁻) mice. Senescence was modulated pharmacologically using the senolytic agent navitoclax (ABT-263), and vascular phenotype was evaluated in adult (6-month-old) and aged mice (18-month-old).

**Results:** PGRN expression increased with age in human and mouse arteries, correlating with elevated p21 expression. PGRN deficiency in adult mice induced endothelial dysfunction, increased vasoconstriction, and induced vascular inflammation and remodeling. Transcriptomic analysis of PGRN⁻/⁻ VSMCs revealed a senescence-associated signature, including perturbed oxidative phosphorylation, altered epigenetic regulation, and collagen pathways. Pharmacological clearance of senescent cells improved endothelial function but increased vascular contractility in PGRN⁻/⁻ mice. In aged mice, PGRN deficiency aggravated vascular dysfunction, remodeling, and renal injury without further increasing senescence markers—suggesting premature, rather than progressive, senescence in the PGRN⁻/⁻ mice.

**Conclusion:** PGRN is a novel regulator of vascular aging, coordinating senescence, inflammation, and remodeling. While endothelial senescence contributes to dysfunction, VSMCs senescence may serve an adaptive role in modulating vascular tone. Targeting PGRN or senescence pathways may offer therapeutic opportunities for age-related vascular diseases, especially in patients with PGRN mutations associated with frontotemporal dementia.

## INTRODUCTION

Aging is a major risk factor for cardiovascular diseases (CVD), which significantly contribute to global mortality^1,2^. As individuals age, arteries experience pathological adaptation that leads to various vascular conditions, including hypertension, atherosclerosis, and coronary artery disease^3^. Strategies to improve the vascular health by targeting age-associated molecular changes can promote healthy aging and extend lifespan. Consequently, strategies aimed at vascular rejuvenation hold geroprotective potential^4,5^. Despite this, the mechanisms driving age-related vascular dysfunction remain poorly understood.

We recently demonstrated that progranulin (PGRN), an anti-inflammatory molecule^6,7^ implicated in certain forms of dementia, including frontotemporal dementia (FTD)^8,9^, plays a crucial role in regulating vascular tone across both small and large vessels^10,11^. Specifically, PGRN deficiency results in increased vasoconstriction in mesenteric arteries^10^ and markedly reduced contractility of aortic vascular smooth muscle cells (VSMCs)^11^. This loss of contractility was associated with impaired mitochondrial quality control and dysregulated autophagy^11^— hallmark features of vascular aging and senescence^12–14^. These findings suggest that PGRN may act as a protective factor against vascular aging. Supporting this premise, Zhu et al. (2020)^15^ showed that cardiac PGRN expression increases with age in mice, and that PGRN deficiency exacerbates age-related cardiac hypertrophy while elevating markers of senescence in the heart. Together, these observations point to an important role for PGRN in modulating age-associated changes in both the vascular and cardiac systems. However, despite emerging evidence of its cardioprotective potential, the role of PGRN in vascular aging and senescence remains unexplored.

In this study, we identified a novel role for PGRN in regulating age-associated vascular dysfunction and senescence. Using human and mouse vascular tissues, along with senescent VSMCs, we demonstrated that PGRN expression increases with age. Importantly, the absence of PGRN resulted in accelerated vascular aging, characterized by enhanced vascular senescence, impaired endothelial function, increased vascular contractility, structural remodeling, and worsened renal damage in aged mice. Mechanistically, we found that PGRN modulates oxidative phosphorylation (OXPHOS) and DNA methylation pathways. Notably, pharmacological clearance of senescent vascular cells improved endothelial function and further increased the vascular contractility in PGRN-deficient mice, suggesting that cellular senescence may represent a hallmark of endothelial dysfunction while also serving a compensatory role in preserving vascular function in the absence of PGRN. Overall, our findings establish PGRN as a critical regulator of vascular aging and highlight a potential dichotomy in senescence, where endothelial senescence contributes to dysfunction, while VSMCs senescence may act adaptively to counteract contractile imbalance. Importantly, our findings suggest clinical concerns for individuals with PGRN mutations—frequently associated with FTD—who may be predisposed to premature vascular dysfunction and end-organ injury due to impaired PGRN signaling.

## METHODS

### Mice

Male and female C57BL/6J wild-type (WT, PGRN^+/+^) and global PGRN knockout (PGRN^−/−^; B6(Cg)-^Grntm1.1Aidi/J^) mice aged 6 and 18 months were used in this study. All animals were maintained on standard chow with ad libitum access to tap water. The 18-month-old WT mice were obtained from the Translational Research Branch, Division of Aging Biology at the National Institute on Aging (NIA), National Institutes of Health (NIH), through a special grant (BRUDER092724SE). Upon arrival at the University of South Alabama vivarium, these mice were allowed a 2–3-week acclimation period prior to experimentation.

All mice were housed in an AAALAC-accredited facility within the College of Medicine at the University of South Alabama. Euthanasia was performed using CO₂ overdose. All procedures were approved by the Institutional Animal Care and Use Committee (IACUC) at the University of South Alabama (protocol #2219557) and conducted in accordance with the Guide for the Care and Use of Laboratory Animals.

### Senolytic Drug Treatment

Six-month-old PGRN^-/-^ mice received intraperitoneal injections of navitoclax (ABT263) at a dose of 50 mg/kg/day for 28 days (2 cycles of 5 days on and 14 days off)^16^. Navitoclax, obtained from Adooq Bioscience-CA-USA, was initially dissolved in dimethyl sulfoxide and then resuspended in corn oil. Six-month-old PGRN^-/-^ mice receiving corn oil were used as a control group. At the time of euthanasia, blood samples were collected, and tissues were harvested and prepared for subsequent analyses.

### Human Coronary Arteries Collection

De-identified human tissues were obtained from cadaveric donor tissues via the Center for Organ Recovery and Education (CORE) under the approval by the University of Pittsburgh Committee for Oversight of Research and Clinical Training Involving Decedents (CORID). A detailed protocol describing tissue collection and handling was published previously^17^. Briefly, coronary arteries were recovered from cadaveric organs and stored in cold Belzer UW Cold Storage Transplant Solution (Bridge to Life) at 4°C for transporting. Coronary arteries were excised and washed with a sterile rinsing solution (sterile PBS supplemented with 2.5 μg/mL of fungicide (Gibco, 15290026), 0.05 mg/mL of gentamicin162 (Gibco, 15710064), and 5 μg/mL of plasmocin (InvivoGen, ant-mpt-1) and maintained at −80°C for immunoblotting.

### Vascular Function

Rings from second-order mesenteric resistance arteries were mounted in a wire myograph (Danysh MyoTechnology) for isometric tension recordings with PowerLab software (AD Instruments) as described before^10,18^. Briefly, rings (2 mm) were placed in tissue baths containing warmed (37 °C), aerated (95% O_2_, 5% CO_2_) Krebs Henseleit Solution (in mmol/L: 130 NaCl, 4.7 KCl, 1.17 MgSO_4_, 0.03 EDTA, 1.6 CaCl_2_, 14.9 NaHCO_3_, 1.18 KH_2_PO_4_, and 5.5 glucose). After 30 minutes of stabilization, curves of tension were performed to adjust the ideal tension for each segment, followed by incubation with potassium chloride (KCl, 120mM). Contractility of mesenteric arteries was tested by cumulative applications of the phenylephrine (PE, 10^−10^–10^−4^ M) and thromboxane A_2_ receptor agonist, U46619 (10^−11^–10^−6^ M). Vasodilation of mesenteric arteries was tested by cumulative applications of the acetylcholine (ACh, 10^−10^–10^−4^ M) and sodium nitroprusside (SNP, 10^−10^–10^−4^ M). A statistical approach was employed to create a curve that depicts the relationship between the concentrations of agonists and the resulting contraction and dilation. This allows for the evaluation of the maximal effect (Emax) and pD2 values for the agonists. The “Emax” refers to the maximum impact achievable by the agonists, and “pD2” represents the negative logarithm of the molar concentration of the agonists required to achieve 50% of the maximum effect (EC50).

### Western Blot

Mesenteric arteries from mice, as well as in coronary artery samples from adult and aged humans were extracted using radioimmunoprecipitation assay buffer (RIPA) buffer (30 mM HEPES, pH 7.4,150 mM NaCl, 1% Nonidet P-40, 0.5% sodium deoxycholate, 0.1% sodium dodecyl sulfate, 5 mM EDTA, 1 mM NaV04, 50 mM NaF, 1 mM PMSF, 10% pepstatin A, 10 μg/mL leupeptin, and 10 μg/mL aprotinin). Total protein extracts were centrifuged at 15,000 rpm/10 min and the pellet was discarded. Proteins from homogenates of mesenteric arteries and kidney (20 μg) were used. Proteins were separated by electrophoresis on a polyacrylamide gradient gel (BioRad Hercules) and transferred to Immobilon-P poly (vinylidene fluoride) membranes. Non-specific binding sites were blocked with 5% skim milk or 1% bovine serum albumin (BSA) in tris-buffered saline solution with tween for 1h at 24 °C. Membranes were then incubated with specific antibodies overnight at 4 °C (Supplementary table 1). After incubation with secondary antibodies, the enhanced chemiluminescence luminol reagent (SuperSignal™ West Femto Maximum Sensitivity Substrate, Thermo-Scientific #34095, Massachusetts, USA) was used for antibody detection.

### Real-Time Polymerase Chain Reaction (qPCR)

mRNA from mesenteric arteries, aorta and kidney were extracted using RNeasy Mini Kit (Quiagen #74106, North Rhine-Westphalia, GER). Complementary DNA (cDNA) was generated by reverse transcription polymerase chain reaction (qPCR) with MultiScribe Reverse Transcriptase (Thermo-Scientific #4319983, Massachusetts, USA). Reverse transcription was performed at 58 °C for 50 min; the enzyme was heat inactivated at 85 °C for 5 min, and real-time quantitative RT-PCR was performed with the PowerTrack™ SYBR Green Master Mix (Thermo-Scientific #A46109, Massachusetts, USA). Sequences of genes as listed in Supplementary Table 2. Experiments were performed in a CFX Opus 384 Real-Time PCR System (Bio-Rad Laboratories, Inc.). Data were quantified by 2ΔΔ Ct and are presented by fold changes versus the control group. Fold change is an indicative of either upregulation or downregulation.

### Histology

The aorta and kidney from PGRN^+/+^ and PGRN^-/-^ mice were collected and placed in a 4% paraformaldehyde (PFA) solution for histologic analysis. After 12 hours in PFA, tissues were placed in 70% ethanol until the day of preparing the samples for histology. Aorta and kidney were embedded in paraffin, then samples were sectioned (10µm) and stained with Masson’s trichrome to analyze the remodeling and structure. Aortic cross-sectional area (CSA) was quantified using the Echo microscope software. Images were acquired in a Keyence microscope (BZ-X series).

### Vascular Smooth Muscle Cells Isolation

We conducted the isolation of VSMCs from the aortas of male PGRN^+/+^ and PGRN^-/-^mice^11^. The isolation procedure followed a well-established enzymatic dissociation protocol. Following the isolation, the VSMCs were cultured in DMEM from Invitrogen Life Technologies. To maintain cell health and preserve their physiological characteristics, the culture medium was supplemented with 10% fetal bovine serum (FBS) obtained from HyClone, along with 100 U/ml penicillin, 100 μg/ml streptomycin, and 10 mmol/L Hepes (pH 7.4) from Sigma-Aldrich. To ensure the viability and functionality of the arterial smooth muscle cells during experimentation, we utilized cells within passages 4 to 8.

### Reactive Oxygen Species Quantification Via Lucigenin-Derived Chemiluminescence Assay

As described before^19^, VSMCs were washed with PBS 1x and harvested in 70-μL lysis buffer (2×10-2 M KH2PO4; 10^-3^ M EGTA, and protease inhibitors: 1μg/mL of aprotinin, 1 μg/mL of leupeptin, and 1 μg/mL of pep-statin). About 50 μL of the sample was added to 175-μL assay buffer (50 mM KH2PO4, 1 mM EGTA, and 150 mM sucrose, pH 7.4 and 5×10^-6^ M lucigenin). Then, the first reading was performed and considered as basal reading. Nicotinamide adenine dinucleotide phosphate [NADPH (10^-4^ M)] was added to each sample, and the luminescence signal was measured, for 30 cycles of 30 s each, in a FlexSation 3 microplate reader (Molecular Devices, San Jose). Basal buffer readings were subtracted from the respective sample readings.

### Cell Cycle Analyzes

Cell cycle arrest was analyzed in PGRN^+/+^ and PGRN^-/-^ VSMCs was assessed using the Cell Cycle Phase Determination Kit, following the manufacturer’s instructions (Cayman, MI, USA).

### Senescence-associated β-galactosidase (SAβG) staining

SAβG staining in PGRN^+/+^ and PGRN^-/-^ VSMCs was performed using a Senescence Detection Kit (MilliporeSigma, KAA002)^20^. VSMCs cultured in 12-well plates were washed twice with PBS and fixed for 10 minutes using the diluted fixation solution provided in the kit, following the manufacturer’s instructions. After fixation, cells were washed three times with PBS, and 1 mL of freshly prepared SAβG staining solution was added to each well. Plates were then wrapped in aluminum foil to protect from light and incubated at 37°C for 24 hours. Following incubation, cells were washed three times with PBS, and images were captured using an Echo microscope.

### Senescence Induction In VSMCs

To evaluate PGRN gene expression in aged VSMCs, cells were exposed to 10 Gy of gamma irradiation^21^ and analyzed five days post-irradiation. Expression of p21 gene expression was assessed to confirm the senescent/aged VSMCs phenotype.

### Sequencing and Gene Count Generation

Total mRNA was isolated from PGRN^+/+^ and PGRN^-/-^ VSMCs using the RNeasy Mini Kit (Qiagen #74106, North Rhine-Westphalia, Germany). RNA quality and concentration were assessed using a NanoDrop spectrophotometer. Samples were then shipped on dry ice via overnight delivery to Novogene (Sacramento, CA, USA) for sequencing.

Sequencing of triplicate samples from PGRN^+/+^ and PGRN^-/-^ VSMCs was performed by Novogene on a fee-for-service basis. Briefly, poly-T oligo magnetic beads were used to isolate messenger RNA (mRNA) from total RNA, which was then fragmented to lengths of 150–350 bases. The mRNA was reverse-transcribed to generate a cDNA library, which was quantified using a Qubit fluorometer, and the size distribution was verified with a Bioanalyzer. The libraries were pooled and sequenced on an Illumina platform to generate paired-end reads. Adapter sequences were removed from the raw reads using Perl scripts. Clean paired-end reads were aligned to the mouse reference genome using HISAT2 (v2.0.5), and gene counts were generated with featureCounts (v1.5.0).

### Transcriptome Data Processing, Differential Expression and Pathway Analysis

Raw gene counts were imported into R (v4.3.1) and analyzed using the standard DESeq2 (v1.42.0) workflow, with samples labeled by genotype: wild type (WT; n = 3) and progranulin-knockout (KO; n = 3). Based on principal component analysis, one KO replicate was excluded from downstream analysis. Normalized counts were obtained using size-factor normalization and row-scaled prior to heatmap generation. For qualitative visualization, the depicted heatmaps display two WT replicates matched to the two KO samples, although all WT samples were included in differential expression testing.

Genes with a Benjamini–Hochberg adjusted p-value (FDR) < 0.05 were considered differentially expressed (DEGs). Senescence-associated genes were identified by intersecting DEGs with Reactome pathways annotated for senescence (retrieved via the msigdbr package (v7.5)). Overlapping genes were grouped into functional categories for visualization. To characterize transcriptional signatures related to DNA methylation, Reactome gene sets corresponding to known regulators of epigenetic homeostasis were retrieved and consolidated into groups based on annotated biological mechanisms. DEGs were intersected with these pathways, excluding genes already included in the senescence panel to avoid redundancy.

### Analysis of Differential DNA Methylation in Mesenteric Arteries

DNA was extracted from mesenteric beds of 6-month-old PGRN^+/+^ (n = 4) and PGRN^-/-^(n =3) using the DNeasy Mini Kit (Qiagen, Germantown, MD). Extracted DNA was then fragmented using g-TUBEs (Covaris, Woburn MA) and 2 µg of sheared DNA was prepared for long-read Nanopore sequencing using the Ligation Sequencing Kit V14 (Oxford Nanopore Technologies, Oxford, UK). Samples were loaded onto individual MinION flow cells; to optimize output, the flow cells were washed (and 150 ng of library re-loaded) twice. DNA methylation was then assessed using Reduced Representation Methylation Sequencing (RRMS) which adaptively enriches for genomic sequences harboring annotated CpG regions.

The resulting reads were basecalled with dorado (v.1.3.0) using a model trained to identify 5mC/5hmC modifications. The reads were aligned to the mm10 mouse reference genome, and the resulting alignment files containing the modified-base information were merged across replicates with samtools (v.1.22.1) to equalize coverage. For each sample, modkit (v.0.5.0) was used to compute 5mC/5hmC frequencies for all genomic CpG positions and assess differences in methylation between the wild-type and knockout genotypes. Individual CpG sites, promoters (defined as −5 kb to +500 bp relative to the TSS), and CpG islands (defined as ±4 kb around island centers) were analyzed for differential methylation, with the resulting summary files filtered for coverage (≥10 reads in each condition) and confidence (≥ 0.8 lower bound of the computed Cohen’s *h* statistic).

High-confidence differentially methylated CpGs were annotated using known mm10 genomic features and custom enhancer, open-chromatin, and CTCF-binding elements from the SCREEN database. To visualize differential methylation patterns across the identified promoters and CpG islands, each region was divided into fixed 50 bp windows and 5mC/5hmC fractions were calculated as the coverage-weighted ratio of modified to total reads. These values were converted into bigWig signal tracks using bedtools (v2.28.0) and supplied to deepTools (v3.5.6) for heatmap generation.

### Statistical Analysis

For comparisons of time-course experiment, one-way analysis of variance (ANOVA) followed by the Tukey post-test was used. Group comparisons were performed using Student’s t test to assess differences between PGRN⁺/⁺ and PGRN⁻/⁻ mice at 6 and 18 months of age, as well as to compare age-related changes within each genotype (6- vs. 18-month-old PGRN⁺/⁺ or PGRN⁻/⁻ mice).

Concentration–response curves were analyzed using a nonlinear interactive fitting program, creating a curve that depicts the relationship between the concentrations of agonists and the resulting contraction/relaxation. This allows for the evaluation of the maximal effect [Emax, refers to the maximum effect achievable by potassium chloride (KCl)] and pD2 values [represents the negative logarithm of the molar concentration of agonist required to achieve 50% of the maximum effect (EC50)].

All analyzes were performed using GraphPad Prism 8.0. (GraphPad Software Inc., San Diego, CA). The level of significance for all variables was 5%.

## RESULTS

### Vascular aging is associated with increased vascular PGRN

First, we assessed PGRN expression in the aged vasculature. PGRN levels were markedly increased in mesenteric and coronary arteries from both aged mice and humans (Figure 1A–B), coinciding with elevated expression of p21, a well-established marker of cellular senescence. The specificity of the PGRN antibody was confirmed by the absence of signal in arteries from PGRN⁻/⁻ mice (Figure 1A). Moreover, a time-course analysis of mesenteric arteries from mice aged 4 to 12 months revealed a progressive rise in PGRN expression that closely paralleled the increase in p21 (Figure 1C). This elevation in PGRN was not influenced by sex, as aged female mice and humans showed similar increases. In addition, VSMCs exposed to irradiation—a known inducer of senescence—also displayed upregulation of both PGRN and p21 (Figure 1D). Together, these findings strongly suggest that PGRN is associated with vascular aging. However, whether this increase represents a protective or maladaptive response remains to be determined.

**Figure 1.**
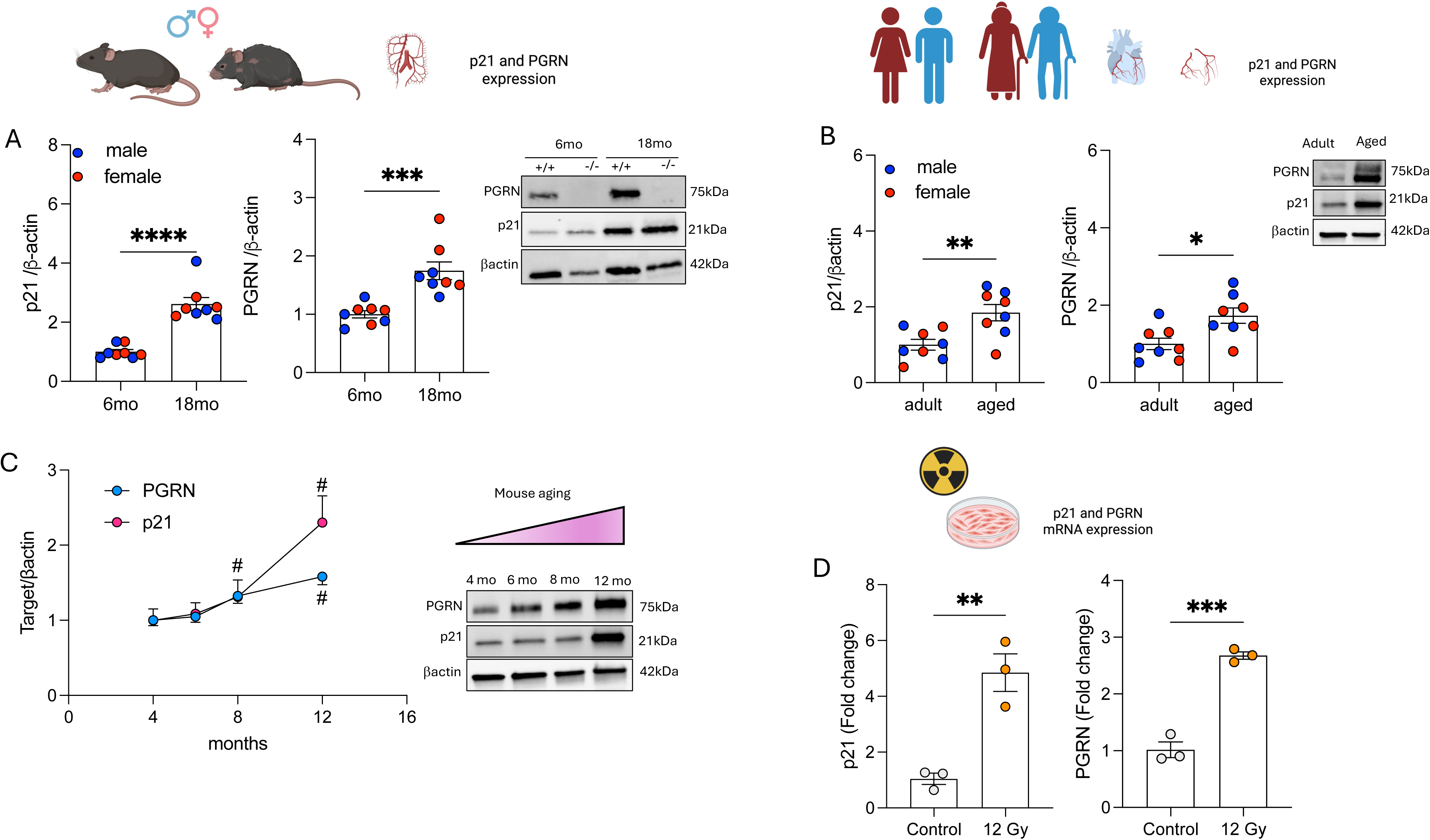
Vascular aging is associated with progranulin (PGRN) expression. (A) p21 and PGRN protein expression in mesenteric arteries from male and female mice at 6 months (adult) and 18 months (aged). (B) p21 and PGRN protein expression in coronary arteries from male and female adult and aged human subjects. (C) p21 and PGRN protein expression in mesenteric arteries from male and female mice aged 4–12 months. (D) p21 and PGRN mRNA expression in vascular smooth muscle cells (VSMCs) exposed or not to irradiation (12 Gy). Data are shown as mean ± SEM, N = 3–8. *P < 0.05 vs. 6-month mice, adult subjects, or control cells. ^#^P < 0.05 vs. 4- and 6-month mesenteric arteries.

### Deficiency in PGRN induces vascular senescence

To explore the role of PGRN in VSMCs biology, we performed RNA sequencing on VSMCs isolated from PGRN+/+ and PGRN-/- mice. Loss of PGRN resulted in substantial transcriptomic changes, with over 2,000 genes differentially expressed (Figure 2A). To determine whether these changes relate to vascular aging, we selected a curated panel of senescence-related genes and grouped them into functional categories. Notably, several of our significant genes mapped to these pathways, including those regulating cell cycle arrest, ubiquitination, DNA damage response, and SASP (senescence-associated secretory phenotype) (Figure 2B). Genes involved in epigenetic regulation were also identified, suggesting that PGRN deficiency perturbs nuclear organization beyond conventional transcriptional control. In line with this, the differential expression of multiple histone genes pointed to potential alterations in chromatin structure, prompting us to examine whether loss of PGRN also impacts DNA methylation, an epigenetic process tightly coupled to histone dynamics. Nanopore-based 5mC profiling revealed a global reduction in DNA methylation, particularly across promoters, CpG islands, and enhancers, consistent with broad epigenomic remodeling (Figure 2C-D) and indicative of a hypomethylated vascular epigenome — a hallmark of vascular aging and senescence — in the absence of PGRN. To further probe whether the transcriptional consequences of PGRN loss could help explain the widespread hypomethylation we observed, we compared our differentially expressed genes to pathways broadly linked to epigenetic stability. These included functional processes that maintain methylation fidelity such as the DNA methylation machinery itself as well as metabolic cofactors, redox regulators, mitochondrial stability, and inflammatory signaling networks — all of which are known modulators of DNA methylation states during cellular senescence. Our results found coordinated dysregulation across these pathways (Figure 2E), suggesting a cellular environment that is less capable of sustaining DNA methylation and providing a plausible transcriptional basis for the hypomethylation observed in PGRN-deficient VSMCs.

**Figure 2.**
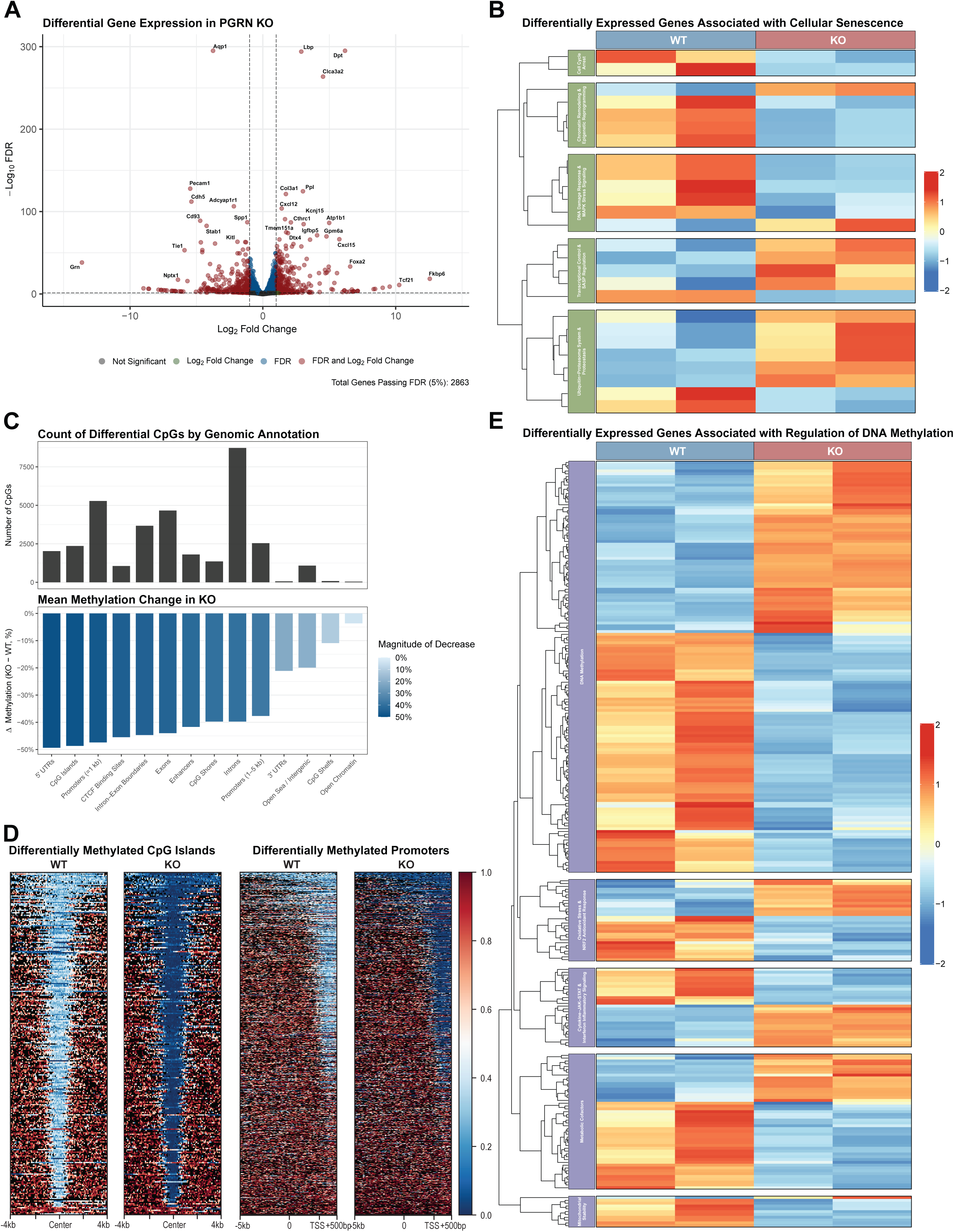
Progranulin (PGRN) deficiency alters vascular transcriptome and epigenetic signatures. (A) Volcano plot of differentially expressed genes (DEGs) in vascular smooth muscle cells (VSMCs) from PGRN+/+ (WT) and PGRN−/− (KO) mice. Genes passing |log₂FC| > 1 and FDR < 0.05 are highlighted, with significance and fold-change thresholds indicated by color. (B, E) Heatmaps showing normalized expression of significant (FDR < 0.05) DEGs that intersected with annotated senescence-related pathways (B) and pathways involved in regulating DNA methylation (E). Columns represent individual WT (blue) and KO (red) replicates; row splits denote functional categories. Colors are reflective of normalized z-scores (−2 to 2), with red indicating higher relative expression and blue indicating lower relative expression. (C) Stacked bar plots summarizing differentially methylated CpG sites between WT and KO mesenteric arteries. Top: Number of high-effect (lower bound of Cohen’s *h* > 0.8) CpGs assigned to each labeled regulatory category. Bottom: Mean methylation differences (KO − WT, percentage points) for the same categories. Bar color reflects the magnitude of methylation loss in KO, with darker shades indicating larger decreases. (D) Heatmaps depicting the average fraction of modified cytosines (5mC + 5hmC) in mesenteric arteries obtained from wild-type (WT) and progranulin-deficient (KO) mice. Signal is plotted over CpG islands (left) and promoters (right) found to be differentially methylated between the two conditions, sorted by average KO signal. Each row represents an individual promoter or CpG island identified as differentially methylated. For CpG islands, the plotted region is ±4 kb flanking the island center; for promoters the plotted region is −5 kb to +0.5 kb relative to the transcription start site (TSS).

The senescent phenotype was further validated in mesenteric arteries from 6-month-old PGRN-/- mice, which showed elevated p21 expression (Figure 3A), and in VSMCs, where we observed increased SaβG activity and impaired cell cycle progression, with PGRN-/- cells exhibiting G1-phase arrest—a hallmark of cellular senescence (Figures 3B and 3C). In line with our previous work identifying PGRN as a key regulator of mitochondrial quality in VSMCs^11^, transcriptomic analysis revealed significant enrichment of genes linked to impaired oxidative phosphorylation in PGRN-deficient cells (Figure 3D). Because mitochondrial dysfunction in oxidative phosphorylation is often coupled to oxidative stress, we next assessed ROS levels. PGRN deletion markedly elevated ROS production in VSMCs, an effect further amplified by antimycin A, a Complex III inhibitor that induces mitochondrial ROS by collapsing the proton gradient and promoting electron leakage (Figure 3E).

**Figure 3.**
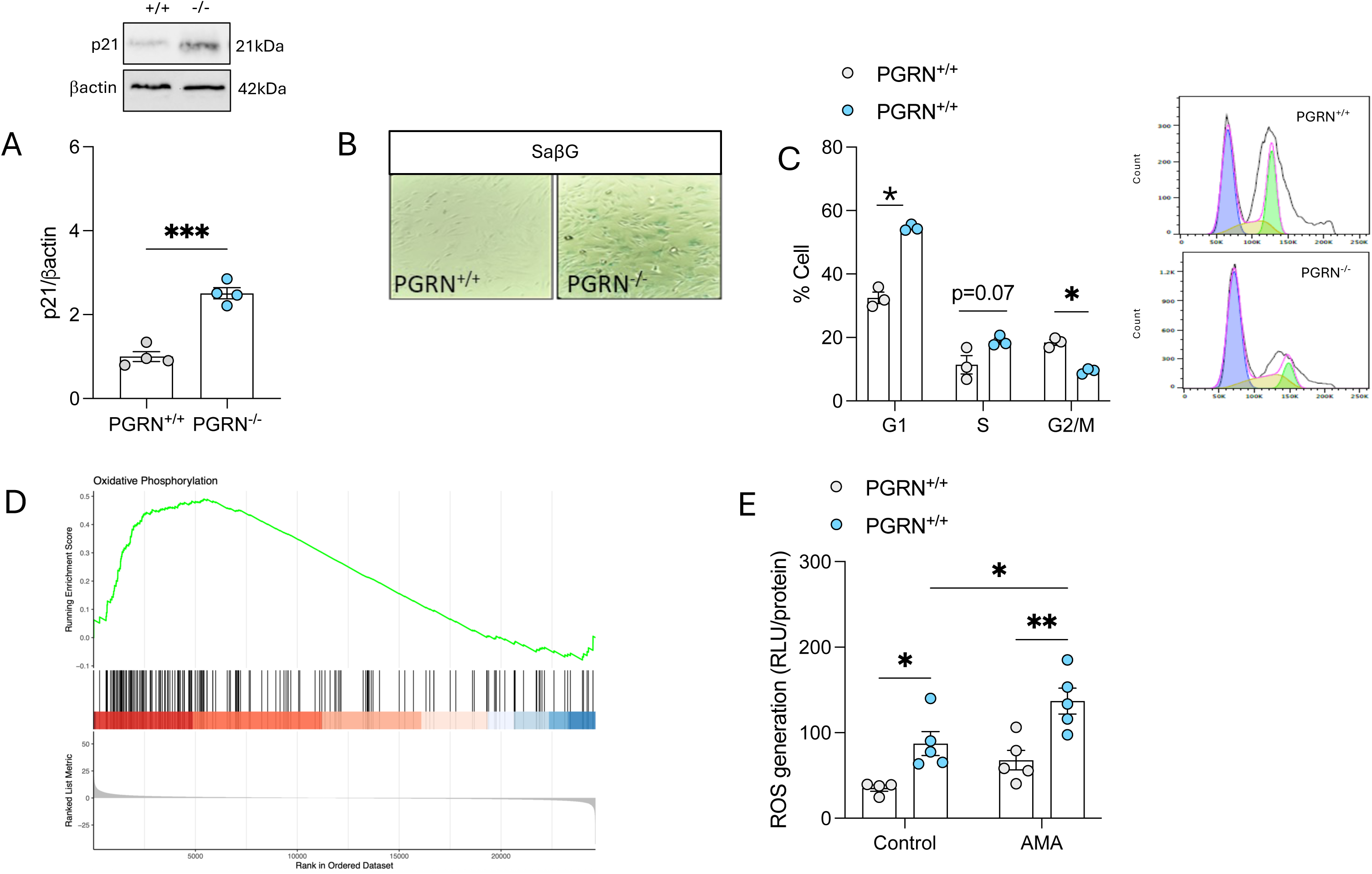
Progranulin (PGRN) deficiency induces premature vascular senescence. (A) p21 protein expression in mesenteric arteries from 6-month-old PGRN+/+ and PGRN-/- mice. (B) Senescence-associated β-galactosidase (SA-β-gal) accumulation in vascular smooth muscle cells (VSMCs) from PGRN+/+ and PGRN-/- mice. (C) Cell cycle analysis of VSMCs from PGRN+/+ and PGRN-/- mice. (D) Gene Set Enrichment Analysis (GSEA) of the oxidative phosphorylation pathway in VSMCs from PGRN+/+ and PGRN-/- mice. (E) Reactive oxygen species (ROS) levels measured by lucigenin in VSMCs from PGRN+/+ and PGRN-/- mice with or without antimycin A treatment (AMA, 20 μM, 30 min). Data are presented as mean ± SEM, N = 3–4. *P < 0.05 vs. PGRN+/+.

### Deficiency of PGRN induces vascular dysfunction and a mild vascular inflammation in adult mice

Primarily we analyzed whether lack of PGRN can affect the vascular function, inflammation, and remodeling, as well as renal fibrosis. We found that deficiency in PGRN does not affect the KCl-induced vascular contraction, but it did increase the vascular contractility for PE and TXA2 mimetic (Figures 4A). It further induced endothelial dysfunction, characterized by an impaired vasodilation response to ACh, without affecting endothelial Nitric Oxide Synthase (eNOS) expression (Supplementary Figure 1A), but did not affect the endothelium-independent vasodilation (SNP response) (Figure 1A). Next, we analyzed markers of vascular phenotype and inflammation in mesentery, although lack of PGRN did not affect α smooth muscle actin (αSMA) and collagen 3α1 (Coll3α1) gene expressions, it did increase Intercellular Adhesion Molecule 1 (ICAM) and interleukin-1 (IL1β) expression. No difference was observed for tumor necrosis factor-α (TNF-α) and Vascular Adhesion Molecule 1 (VCAM) (Figure 4B).

**Figure 4.**
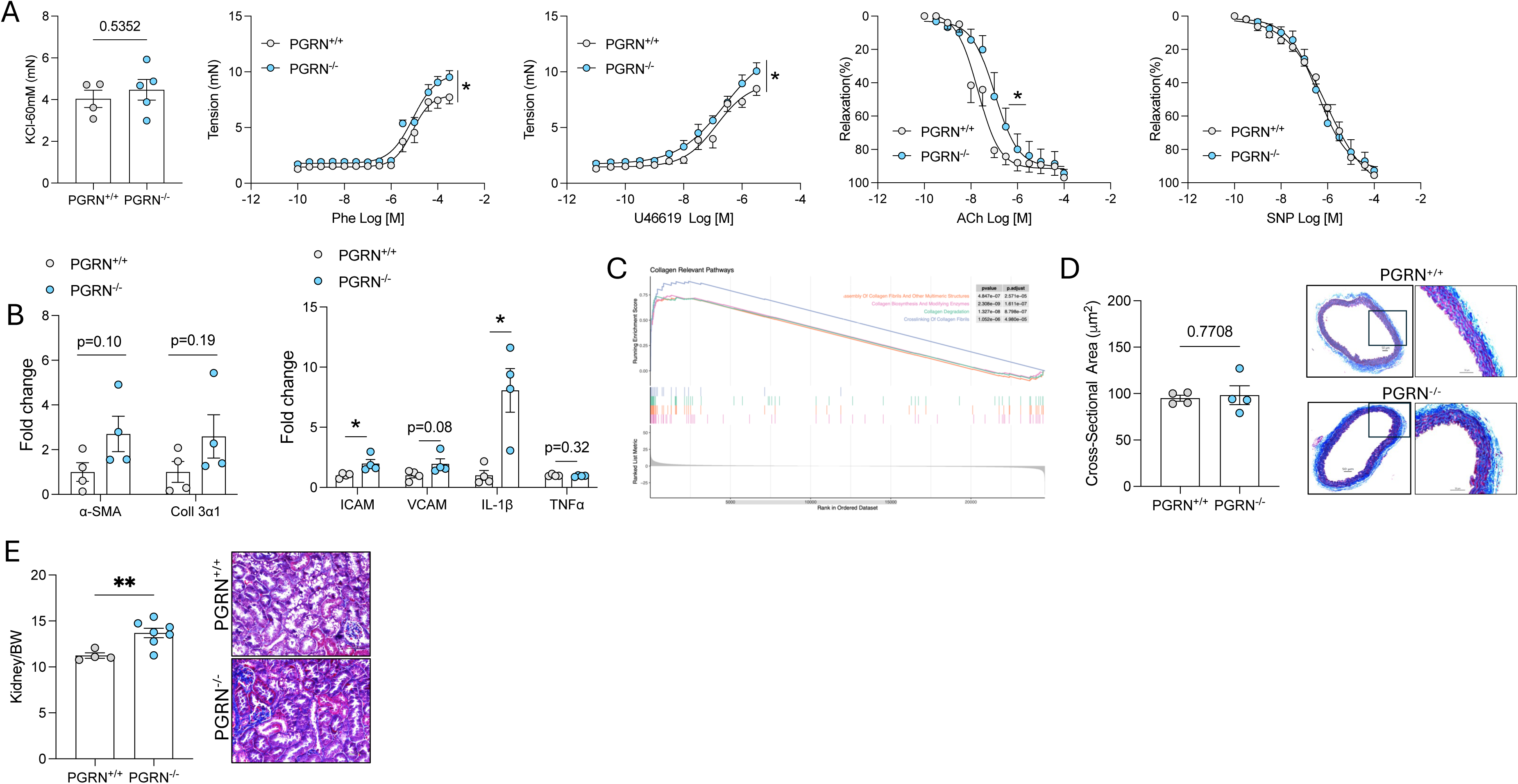
Progranulin (PGRN) deficiency impairs vascular function and promotes inflammation in mesenteric arteries. (A) KCl-induced contraction and concentration–response curves (CRCs) to phenylephrine, U46619, acetylcholine (ACh), and sodium nitroprusside (SNP) in mesenteric arteries from 6-month-old PGRN+/+ and PGRN-/-mice. (B) Expression of structural and inflammatory genes in mesenteric arteries from 6-month-old PGRN+/+ and PGRN-/- mice. (C) Gene Set Enrichment Analysis (GSEA) highlighting collagen-related pathways in VSMCs from PGRN+/+ and PGRN-/- mice. (D) Aortic cross-sectional area and representative Masson’s Trichrome staining of aortas from 6-month-old PGRN+/+ and PGRN-/- mice. (E) Kidney weight and representative Masson’s Trichrome staining of kidneys from 6-month-old PGRN+/+ and PGRN-/- mice. Data are shown as mean ± SEM, N = 4-7. *P < 0.05 vs. PGRN+/+.

To assess end-organ injury, we examined whether the absence of PGRN in adulthood affects the aorta and kidney, two key targets of aging. Transcriptomic analysis of VSMCs from PGRN+/+ and PGRN-/- mice revealed strong enrichment of collagen-related pathways, including collagen fibril assembly, biosynthesis, modification, degradation, and crosslinking (Figure 4C). At 6 months of age, PGRN-/- mice did not exhibit overt aortic thickness; however, they showed increased collagen deposition (Figure 4D), elevated αSMA expression, and enhanced vascular inflammation, evidenced by upregulation of VCAM, IL-1β, and TNF-α (Supplementary Figure 1B and C), suggesting that the absence of PGRN may promote vascular inflammation and fibrosis through dysregulated extracellular matrix dynamics.

Furthermore, PGRN-/- mice exhibited increased renal mass (Figure 4E), elevated fibrotic markers (Figure 4E) with no effects on renal inflammation (Supplementary Figure 1D and E), and higher expression of neutrophil gelatinase-associated lipocalin (NGAL), a marker of kidney disease progression (Figure 5F–I).

**Figure 5.**
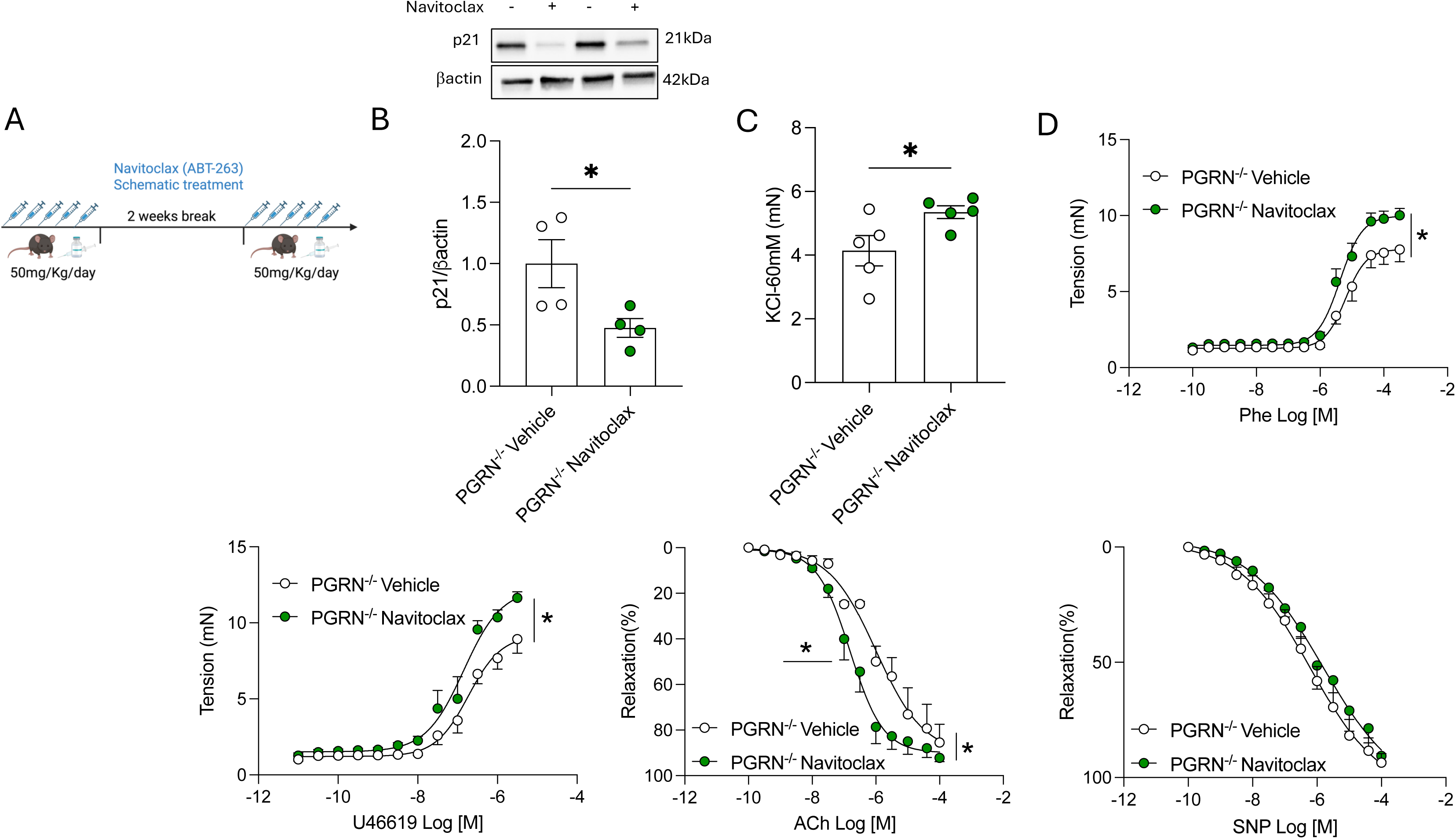
Senolytic treatment modulates vascular function in progranulin (PGRN)–deficient mice. (A) Schematic of Navitoclax (ABT-263, 50 mg/kg/day) treatment. (B) p21 protein expression. (C) KCl-induced contractility. (D) Concentration–response curves (CRCs) to phenylephrine, U46619, acetylcholine (ACh), and sodium nitroprusside (SNP). All experiments were performed in mesenteric arteries from PGRN-/- mice treated with vehicle (5% DMSO, 95% corn oil) or Navitoclax. Data are shown as mean ± SEM, N = 4. *P < 0.05 vs. PGRN-/- vehicle.

### Elimination of vascular senescent cells improves endothelial function but exacerbates vascular contraction of mesenteric arteries

To investigate the contribution of vascular senescence to vascular dysfunction, we treated 6-month-old PGRN^-/-^ mice with the senolytic drug navitoclax (ABT-263) (Figure 5A). This treatment effectively reduced vascular senescence in PGRN-deficient mice characterized by a significant decrease in mesenteric p21 (Figure 5B). Interestingly, despite this reduction, navitoclax further enhanced vascular contractile responses to KCl (Figure 5C), PE, and U46619 (Figure 5D). Notably, navitoclax treatment restored endothelial function, as evidenced by improved vasodilation, without altering responses to the endothelium-independent vasodilator SNP (Figure 5D).

### Deficiency of PGRN exacerbates aging-induced vascular injury

As expected, aged male PGRN^+/+^ mice (18 months old) displayed marked vascular and endothelial dysfunction, accompanied by significant changes in inflammatory, remodeling, and senescence markers in the mesenteric arteries, aorta, and kidneys, compared to adult male PGRN^+/+^ mice (6 months old) (Supplementary figures 2 and 3). To determine whether exacerbated senescence could further worsen the vascular phenotype in the absence of PGRN, we compared 18-month-old PGRN^+/+^ and PGRN^-/-^mice. While PGRN deficiency did not alter KCl-induced vascular contractility (Figure 6A), it significantly enhanced vasoconstrictive responses to PE and U46619 (Figure 6A). In addition, PGRN^-/-^ mice displayed aggravated endothelial dysfunction (Figure 6A) followed by increased eNOS gene expression (Supplementary figure 4A), and impaired vasodilation in response to SNP (Figure 6A).

**Figure 6.**
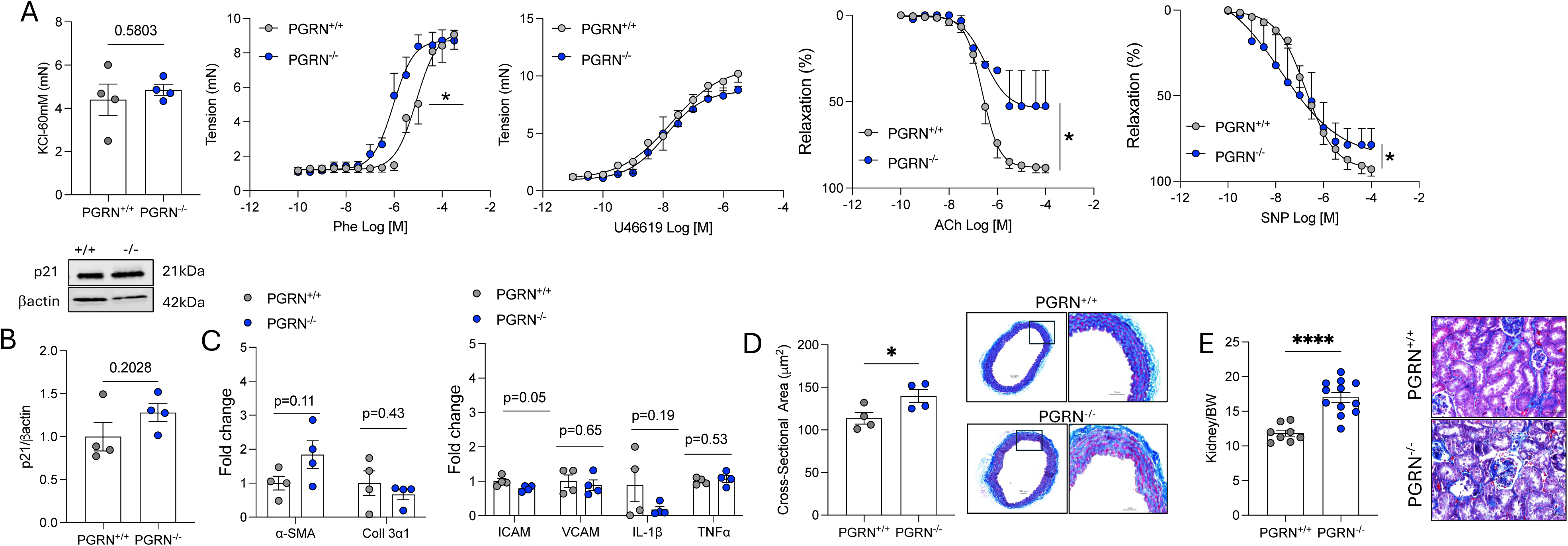
Progranulin (PGRN) deficiency does not further increase vascular senescence but aggravates age-associated vascular dysfunction. (A) KCl-induced contraction and concentration–response curves (CRCs) to phenylephrine, U46619, acetylcholine (ACh), and sodium nitroprusside (SNP) in mesenteric arteries from 18-month-old PGRN+/+ and PGRN-/- mice. (B) p21 protein expression. (C) Expression of structural and inflammatory genes in mesenteric arteries from 18-month-old PGRN+/+ and PGRN-/- mice. (D) Aortic cross-sectional area and representative Masson’s Trichrome staining of aortas from 18-month-old PGRN+/+ and PGRN-/- mice. (E) Kidney weight and representative Masson’s Trichrome staining of kidneys from 18-month-old PGRN+/+ and PGRN-/- mice. Data are shown as mean ± SEM, N = 4-12. *P < 0.05 vs. PGRN+/+.

Interestingly, vascular senescence levels—indicated by p21 expression—remained unchanged between aged PGRN^+/+^ and PGRN^-/-^ mice (Figure 6B). Our data suggest that PGRN deficiency may advance the onset of vascular senescence (premature senescence) rather than further increasing senescence burden at advanced age; however, this remains inferential given we primarily assessed p21 and a limited SASP profile. Similarly, markers of vascular inflammation and remodeling in the mesenteric arteries were not further altered by PGRN deficiency (Figure 6C).

In the aorta, we also observed no additional changes in inflammatory or structural markers (Supplementary figure 4B and C); however, CSA was notably increased in aged PGRN^-/-^ mice (Figure 6D). Renal injury is a well-recognized hallmark of aging and cellular senescence. Therefore, we examined whether PGRN deficiency could exacerbate kidney damage in aged mice. While no significant differences were observed in inflammatory markers between kidneys of aged PGRN^+/+^ and PGRN^-/-^ mice (Supplementary figure 4D and E), PGRN^-/-^ mice exhibited increased kidney mass (Figure 6D), elevated expression of Kidney Injury Molecule-1 (KIM1), and an expanded fibrotic core (Figure 6D and Supplementary figure 4D and E)—indicating aggravated renal injury in the absence of PGRN.

## DISCUSSION

Our findings identify PGRN as both a marker and regulator of vascular aging. We demonstrate that PGRN expression increases in mouse and human arteries with age, closely paralleling the induction of the senescence marker p21. Loss of PGRN leads to premature activation of vascular senescence pathways, mitochondrial dysfunction, epigenetic remodeling, inflammation, fibrosis, and renal injury—phenotypes that collectively resemble early vascular aging. These observations support a model in which PGRN upregulation represents a compensatory response aimed at preserving vascular integrity during aging.

The increased expression of PGRN in aged arteries and in VSMCs exposed to irradiation suggests that PGRN is engaged during vascular stress and may counter the detrimental effects of senescence. Whether this upregulation is protective, maladaptive, or both likely depends on the stage of vascular aging and the balance between mitochondrial dysfunction, inflammation, and chromatin remodeling. Previous studies from our lab in vascular biology^10,11^ and others in neurodegeneration^9^ and cardiac physiology^15^ have shown that PGRN exerts anti-inflammatory, anti-pathological remodeling, and mitochondria and lysosome-stabilizing effects, whereas its deficiency leads to lysosomal dysfunction and neuronal loss In this context, elevated PGRN during aging may reflect an adaptive attempt to maintain vascular homeostasis under chronic stress, although this compensatory response may become insufficient as aging progresses.

Loss of PGRN in VSMCs triggered profound transcriptomic and epigenomic reprogramming, with more than 2000 genes differentially expressed and extensive remodeling of DNA methylation landscapes. Senescence-associated pathways—including those related to chromatin architecture, ubiquitination, DNA damage response, and SASP—were prominently enriched, indicating that PGRN deficiency perturbs nuclear homeostasis and genomic stability, hallmarks of aging^22,23^. Notably, however, the overlap between differentially expressed genes and those associated with differentially methylated regions was limited, suggesting a temporal dissociation between methylation and transcription, with promoter 5mC depletion possibly “priming” chromatin for later transcriptional activation. Together, these findings indicate that PGRN regulates chromatin state at multiple levels—through histone gene expression and nucleosome dynamics for rapid transcriptional responses, and through DNA methylation to provide long-term epigenetic stability. Histone acetylation and methylation likely mediate the immediate transcriptional reprogramming observed in PGRN deficiency, whereas DNA methylation acts as a secondary stabilizing mechanism. Future work should interrogate histone post-translational modifications to elucidate how PGRN coordinates chromatin remodeling with vascular senescence.

Our transcriptomic data also highlight several mechanisms that may underlie the reduced DNA methylation observed in PGRN-deficient cells. The KO VSMCs exhibited clear evidence of oxidative stress—consistent with our previous work demonstrating mitochondrial-driven ROS disruption—and reduced antioxidant capacity, creating a redox environment unfavorable for maintaining cytosine modifications. Under such conditions, ROS can oxidize 5-methylcytosine and related bases, generating lesions that are removed by base-excision repair and replaced with unmodified cytosines^24^. This repair-mediated cytosine turnover offers a parsimonious explanation for the broad loss of both methylation and hydroxymethylation in PGRN-deficient cells, independent of an active demethylation program. Given that mitochondrial stress is both a cause and amplifier of premature senescence^25^, these redox-driven effects likely link mitochondrial dysfunction to the widespread hypomethylation and accelerated aging phenotype observed in the absence of PGRN.

In parallel, several pathways that support methylation fidelity—including metabolic cofactors, mitochondrial stability, and inflammatory signaling^26,27^—were also disrupted, creating conditions in which both maintenance methylation and TET-dependent cytosine modification are likely impaired. Because cellular senescence is fundamentally an epigenetically remodeled state characterized by hypomethylation^28^, the coordinated dysregulation of these processes in PGRN-deficient cells provides a unifying framework. Although the precise mechanism remains uncertain, the combined transcriptomic and methylation patterns are consistent with a model in which stress-driven cytosine turnover and impaired methylation maintenance together contribute to the widespread CpG hypomethylation observed in the absence of PGRN.

Adult PGRN-/- mice manifested early vascular dysfunction characterized by heightened vasoconstriction to phenylephrine and thromboxane A₂ mimetic, as well as impaired endothelial-dependent relaxation. These changes occurred despite preserved eNOS mRNA expression, suggesting post-transcriptional or functional impairment of endothelial signaling. The upregulation of ICAM-1 and IL-1β, along with enrichment of collagen-related pathways, indicates early inflammatory activation and extracellular matrix dysregulation. Increased collagen deposition and αSMA expression in the aorta further support the role of PGRN in restraining vascular fibrotic remodeling. Concomitant renal enlargement, increased fibrotic markers, and elevated NGAL expression point to early cardiorenal involvement, consistent with the established link between vascular senescence, renal dysfunction, and hypertension-associated end-organ damage^29,30^.

Transcriptomic profiling revealed a robust enrichment of collagen-related pathways in PGRN-deficient VSMCs, and *in vivo*, PGRN⁻/⁻ mice exhibited increased vascular collagen deposition and elevated αSMA expression—hallmarks of augmented fibrotic remodeling. These structural alterations were accompanied by higher expression of VCAM-1, IL-1β, and TNF-α, reinforcing the concept that PGRN is required to restrain vascular inflammation and extracellular matrix dysregulation. Although alterations in PGRN levels are often associated with fibrotic remodeling^31^, the broader literature suggests that PGRN functions predominantly as an anti-fibrotic factor across several organs—including skin^32^, liver^33^, heart^34^, and lung^35^—with effects that remain highly context-dependent. Our findings are consistent with this anti-fibrotic role and extend it to the vasculature. Importantly, this study is the first to show, through an integrated omics approach, that PGRN deficiency directly activates collagen biosynthesis and extracellular matrix pathways in VSMCs, providing new mechanistic insight into how PGRN governs vascular fibrotic remodeling.

End-organ analysis shown increased kidney mass, elevated fibrotic markers, and higher NGAL expression, indicating that vascular inflammation and fibrosis extend their detrimental effects to the kidney. These findings are consistent with prior studies showing that PGRN plays a crucial role in preserving renal phenotype in models of diabetes and ischemia/reperfusion injury^36,37^. This vascular–renal interaction supports the concept that vascular senescence is an important driver of end-organ damage during aging. Although our study did not establish the temporal sequence of these events, our previous work in younger mice^10^ suggests that vascular dysfunction emerges before overt renal injury.

To dissect the contribution of vascular senescence to these phenotypes, we treated PGRN-/- mice with the senolytic agent navitoclax. Senolytic therapy effectively lowered p21 expression and improved endothelial-dependent vasodilation, confirming a detrimental role of endothelial senescence in impaired relaxation. Surprisingly, however, navitoclax also further increased vascular contractility. These divergent effects highlight that senescent cells within the vascular wall do not exert uniform functional consequences. Endothelial senescence appears deleterious for vasodilatory signaling, whereas VSMCs senescence may serve a compensatory or protective role by limiting excessive contractile tone. This concept is consistent with recent evidence showing that senescent VSMCs exhibit reduced mechanical contractility^38^, which may act as an intrinsic brake on vasoconstriction during vascular disease progression. Because navitoclax is not cell-type–specific, it likely eliminated senescent endothelial cells, VSMCs, and potentially perivascular cells, making the net effect a composite of different cellular contributions. This complexity may help explain the mixed vascular outcomes reported with senolytic therapies in both preclinical and clinical studies.

Importantly, navitoclax has known off-target effects, including thrombocytopenia^39^ due to Bcl-xL inhibition. We also did not assess potential navitoclax-induced vascular apoptosis or activation of the caspase-3 pathway^40^, which represents an additional limitation given that senolytic agents can trigger apoptotic clearance of stressed or damaged cells. Although we did not evaluate platelet counts or apoptotic markers systematically, these unmeasured effects should be considered when interpreting the functional consequences of senolytic treatment.

In aged mice, PGRN deficiency aggravated endothelial dysfunction and vascular hypercontractility without further increasing p21 expression or amplifying inflammatory or remodeling markers. This pattern suggests premature, rather than progressive, vascular senescence: PGRN⁻/⁻ mice appear to reach a senescent threshold earlier in life, after which senescence burden does not substantially increase with chronological aging. Our data suggest that PGRN deficiency advances the onset of vascular senescence (premature senescence) rather than further increasing senescence burden at advanced age; however, this remains inferential given we primarily assessed p21 and a limited SASP profile. This trajectory differs from findings in cardiac tissue, where Zhu et al. (2020)^15^ reported that aged PGRN-deficient mice exhibit further accumulation of senescence markers and worsened cardiac hypertrophy. Several factors may account for this discrepancy. Senescence is highly tissue-specific, and the heart and vasculature differ markedly in cellular composition, metabolic demand, and regenerative capacity. Thus, cardiac cells may continue to accrue senescent burden with age, whereas vascular cells—particularly VSMCs—may reach a senescence plateau earlier. Additionally, the prior study assessed a broader panel of senescence markers, while our analysis focused primarily on p21 and selected SASP components, which may capture different dimensions of the senescence program. Distinct inflammatory and mitochondrial stress environments in cardiac versus vascular tissue may further influence how PGRN deficiency shapes aging trajectories. Together, these considerations highlight that PGRN-dependent aging phenotypes are organ-specific and that PGRN loss may drive premature vascular senescence without progressive expansion of senescent burden in late life. Nonetheless, aged PGRN⁻/⁻ mice exhibited increased aortic cross-sectional area and worsened renal injury, indicating that early-life vascular dysfunction predisposes to exacerbated age-related cardiorenal pathology.

It is important to knowledge the limitations of our study, we primarily focused on p21 as a marker of senescence, and while supported by SA-β-gal staining and SASP gene expression, broader senescence characterization (e.g., p16, γH2AX, telomere dysfunction) was not performed. Blood pressure measurements were not included in this study, which limits direct inference regarding hypertension; however, the observed resistance artery hypercontractility, endothelial dysfunction, renal fibrosis, and our previous findings^10^ strongly suggest a pro-hypertensive phenotype. Finally, although our multi-omics approaches revealed substantial epigenetic remodeling, the mechanistic links connecting PGRN signaling, chromatin state, and mitochondrial homeostasis remain to be defined. This is particularly relevant given that we and others have previously demonstrated a strong relationship between PGRN and mitochondrial function, including its roles in mitophagy, mitochondrial quality control, and metabolic regulation^11,37,41^.

In conclusion, our findings establish PGRN as a mechanistic regulator of vascular aging. PGRN deficiency induces premature vascular senescence and disrupts mitochondrial, epigenetic, and inflammatory homeostasis, resulting in early vascular dysfunction, structural remodeling, and renal injury. Over time, these alterations recapitulate the hallmarks of natural vascular aging but occur earlier and with greater severity. Importantly, the dichotomous effects of senolytic therapy reveal that endothelial and VSMCs senescence have distinct functional consequences in vascular physiology. Given the availability of PGRN-targeted therapeutics in clinical development for frontotemporal dementia, these findings raise important considerations for vascular and cardiorenal health in individuals with PGRN insufficiency or mutations. PGRN may therefore serve both as a biomarker and potential therapeutic target for age-related vascular and cardiorenal disease.

### Perspectives

Progranulin has emerged as a key determinant of how the vasculature ages. By combining human and mouse data with functional, transcriptomic, and epigenetic analyses, we show that progranulin deficiency accelerates vascular senescence, disrupts mitochondrial and chromatin homeostasis, and precipitates early resistance artery dysfunction, fibrotic remodeling, and cardiorenal injury. These findings suggest that progranulin upregulation during aging is not merely a biomarker of stress but part of an adaptive program that helps preserve vascular integrity. Our senolytic experiments further indicate that different senescent vascular cell types have distinct functional roles, with endothelial senescence being clearly detrimental while VSMCs senescence may partially restrain excessive vasoconstriction. Looking forward, defining the upstream signals that regulate vascular progranulin expression, the downstream pathways that couple progranulin to mitochondrial and epigenetic remodeling, and the blood pressure consequences of progranulin loss in vivo will be essential. Given that progranulin-targeted therapies are already in development for frontotemporal dementia, our results raise the possibility that modulating progranulin or its signaling network could be leveraged to delay vascular aging and prevent cardiorenal complications in at-risk populations.

### Novelty and Relevance

#### What Is New?

- Identifies progranulin as a regulator of vascular aging and senescence.
- Shows that progranulin deficiency causes premature vascular dysfunction, fibrosis, and renal injury.
- Uses multi-omics to link progranulin loss to mitochondrial and epigenetic remodeling in vascular smooth muscle cells.

#### What Is Relevant?

- Resistance artery hypercontractility and endothelial dysfunction in progranulin-deficient mice support a pro-hypertensive vascular phenotype.
- Findings suggest that low progranulin states may predispose to earlier onset of hypertension-related vascular and renal damage.

#### Clinical/Pathophysiological Implications?

- Individuals with progranulin mutations or reduced progranulin levels (e.g., GRN-associated frontotemporal dementia) may be at increased vascular and cardiorenal risk.
- Targeting progranulin pathways, or selectively modulating senescent vascular cell populations, may offer new strategies to prevent or treat age-related hypertension and its complications.

---

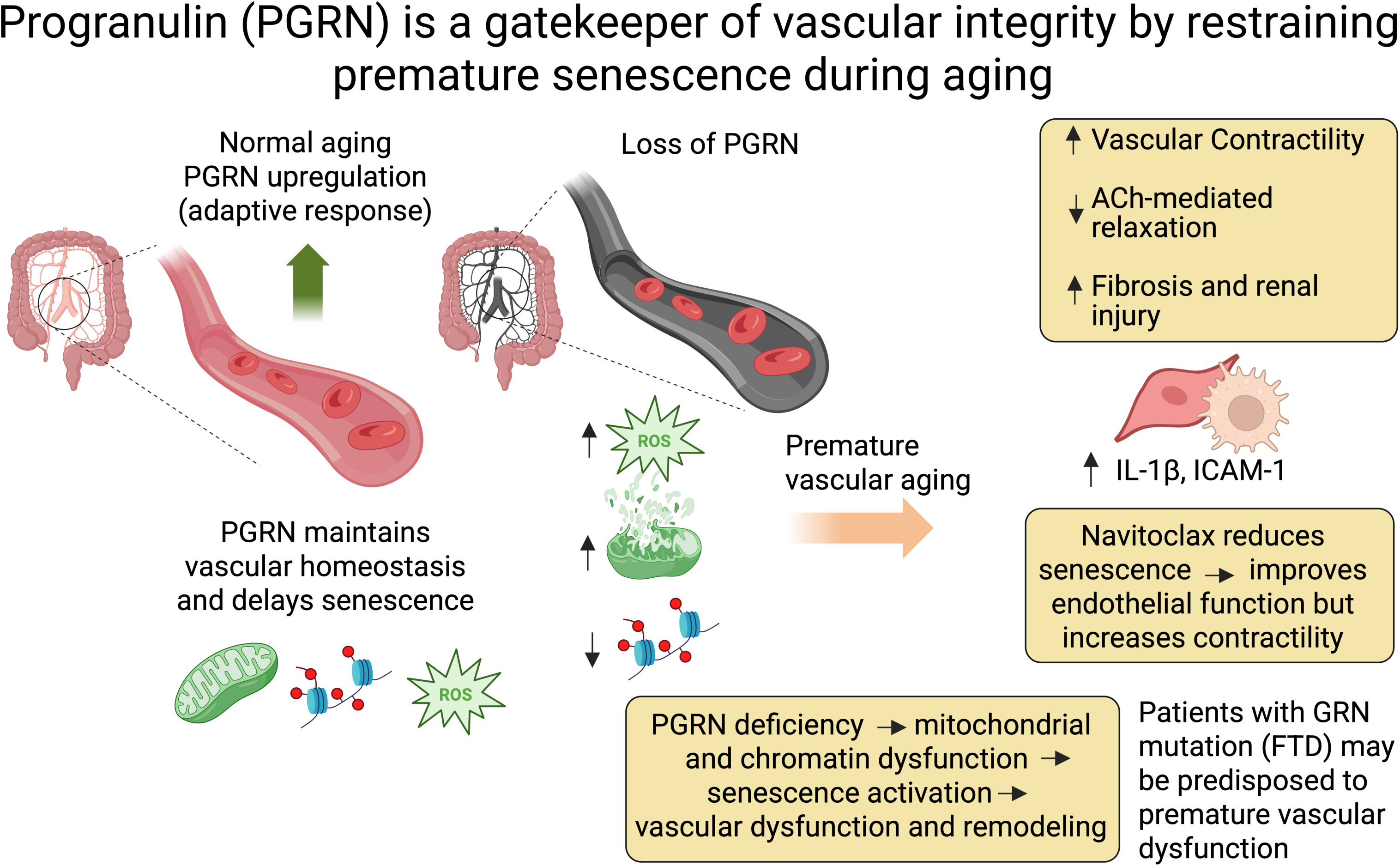

## Notes

### Competing Interest Statement

The authors have declared no competing interest.

### Summary of Updates

Our author's list was not correct. We fixed it. We also added the supplementary files.

### Funding

- NIH Common Fund: HL169202

