## Supplemental files for "Progranulin deficiency aggravates aging-induced vascular injury"

### Material and methods

#### Table S1. Patient information

| Patient ID | Age | Sex | Tissue Type | Cause of Death | Hyper-tension | Diabetes | Smoking | Other Features |
|---|---|---|---|---|---|---|---|---|
| 2016-037 | 31 | Male | LAD Coronary | Anoxia | Unknown | Unknown | Unknown | Dyslipidemia |
| 2016-078 | 34 | Female | LAD Coronary | Heat failure | Unknown | Unknown | Unknown | N/A |
| 2017-069 | 37 | male | LAD Coronary | Accident/Cardiac death | Unknown | Unknown | Unknown | N/A |
| 2019-065 | 39 | Female | LAD Coronary | Overdose | No | No | Unknown | N/A |
| 2016-093 | 43 | Male | LAD Coronary | Cardiac Arrest | Yes | Yes | No | Pneumonia CAD |
| 2019-015 | 45 | Female | LAD Coronary | Cardiac Arrest/Brain Anoxia | Unknown | Yes | Yes | Low BMI |
| 2017-035 | 48 | Female | LAD Coronary | Brain Anoxia | Unknown | Unknown | Unknown | Tumor on intestine, benign |
| 2016-023 | 49 | Female | LAD Coronary | Stroke | Unknown | Unknown | Yes | High BMI |
| 2017-062 | 66 | Female | LAD Coronary | Scleroderma | Unknown | Unknown | Unknown | N/A |
| 2017-042 | 69 | Female | RCA Coronary | Stroke | Unknown | Unknown | Unknown | Pacemaker |
| 2016-060 | 73 | Male | LAD Coronary | Cardiac Arrest | Unknown | Unknown | Unknown | Hypercholesteremia |
| 2016-038 | 81 | Male | LAD Coronary | Heart Attack | Unknown | Unknown | Unknown | N/A |
| 2016-169 | 82 | Female | LAD Coronary | Unknown | Unknown | Unknown | Unknown | N/A |
| 2017-044 | 87 | Female | LAD Coronary | Heart Attack | Unknown | Unknown | 25 years ago | Hyperlipidemia, MI, AVD, COPD |
| 2016-095 | 87 | Male | LAD Coronary | MI | Yes | No | 50 years ago | N/A |
| 2017-046 | 66 | Male | LAD Coronary | Respiratory failure | Unknown | Unknown | Unknown | IPF, PH |

AVD= Aortic Valve Disease, BMI= Body Mass Index, CAD= Coronary Artery Disease, COPD= Chronic Obstructive Pulmonary Disease, IPH= Idiopathic Pulmonary Fibrosis, LAD= Left Anterior Descending Artery, MI= Myocardial Infarction, PH= Pulmonary Hypertension, RCA= Right Coronary Artery.

**Table S2. List of antibodies**

| Antibody | Dilution | Catalog number | Company |
|---|---|---|---|
| p21 | 1:500 | SC6246 | Santa Cruz |
| PGRN (mouse samples) | 1:1000 | AF2557 | R&D |
| PGRN (human samples) | 1:500 | AF2420 | R&D |
| β-actin | 1:20000 | A3854 | Sigma |

**Table S3. List of primers**

| Target genes | | Sequence |
|---|---|---|
| p21 | FW | TCGCTGTCTTGCACTCTGGTGT |
| | RV | CCAATCTGCGCTTGGAGTGATAG |
| PGRN | FW | GTTCCCTGCACAAAAGACCAA |
| | RV | GGGTCTTAGCATCAGGGCAC |
| eNOS | FW | CGCAAGAGGAAGGAGTCTAGCA |
| | RV | TCGAGCAAAGGCACAGAAGTGG |
| IL1b | FW | TGACGGACCCCAAAAGATGA |
| | RV | GCTCTTGTTGATGTGCTGCT |
| aSMA | FW | TGCTGACAGAGGCACCACTGAA |
| | RV | CAGTTGTACGTCCAGAGGCATAG |
| Collagen 3a1 | FW | CTGAAGATGTCGTTGATGTG |
| | RV | ACTGTCTTGCTCCATTCC |
| ICAM | FW | ATCACATGGGTCGAGGGTTT |
| | RV | AACCACTGCCAGTCCACATA |
| VCAM | FW | TGACAAGTCCCCATCGTTGA |
| | RV | ACCTCGCGACGGCATAATT |
| TNFa | FW | AATGGCCTCCCTCTCATCAG |
| | RV | CCTAACTGCCCTTCCTCCAT |
| Podocin | FW | GACACCTATCACAAGGTTGA |
| | RV | CCATGCGGTAGTAGCAGACA |
| Lcn2 (NGAL) | FW | TGGCCCTGAGTGTCATGTG |
| | RV | CTCTTGTAGCTCATAGATGGTGC |
| Havcr1 (KIM-1) | FW | ACATATCGTGGAATCACAACGAC |
| | RV | ACAAGCAGAAGATGGGCATTG |
| GAPDH | FW | GAGAGGCCCTATCCCAACTC |
| | RV | TCAAGAGAGTAGGGAGGGCT |

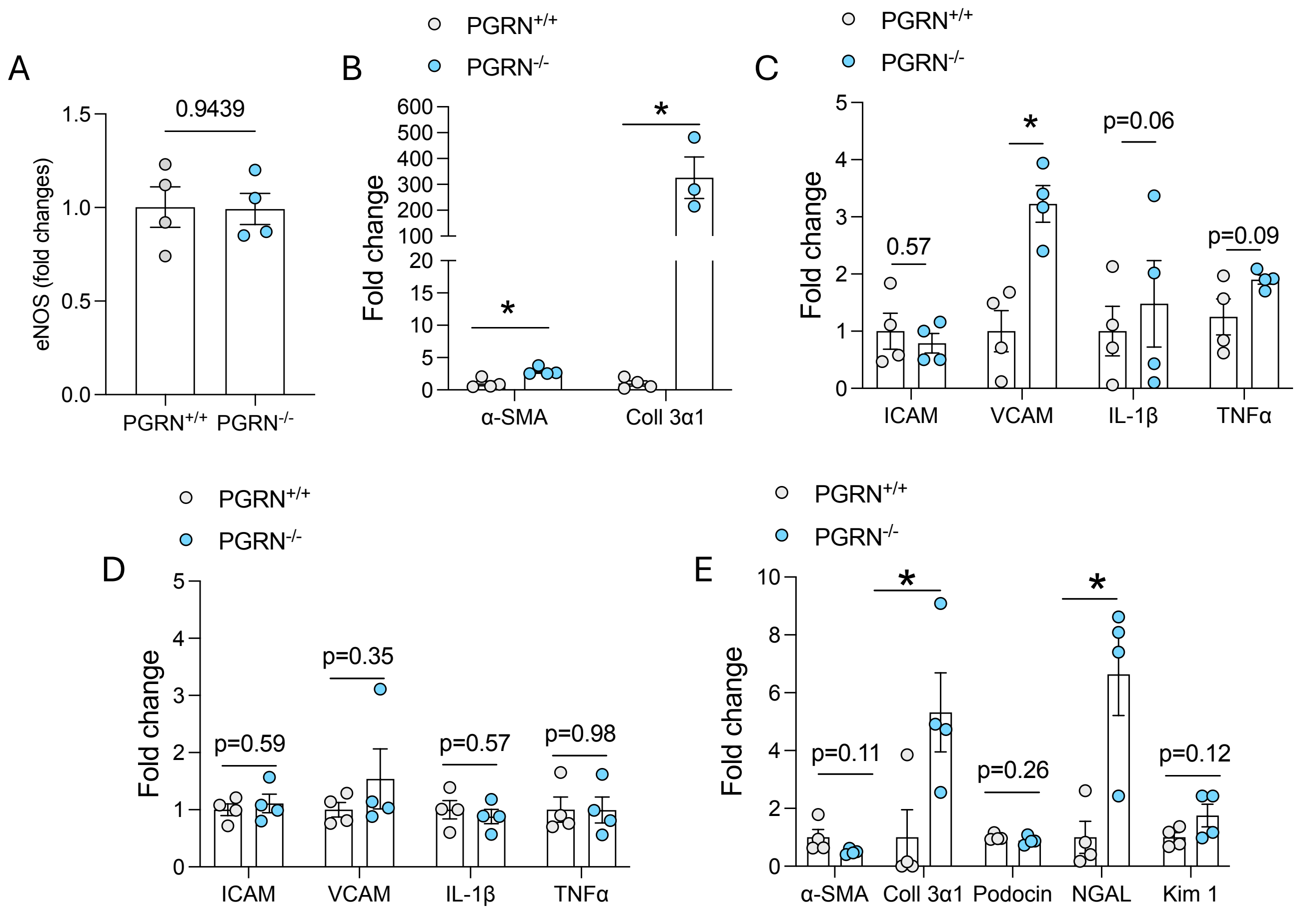


**Supplementary Figure 1. Progranulin (PGRN) deficiency promotes injury in mesenteric arteries end-organ damage.** (A) eNOS mRNA expression in mesenteric arteries. (B–C) Expression of structural and inflammatory genes in aorta. (D–E) Expression of structural and inflammatory genes in kidneys. All experiments were conducted in samples from 6-month-old PGRN+/+ and PGRN-/- mice. Data are shown as mean ± SEM, N = 4-5. *P < 0.05 vs. PGRN+/+.

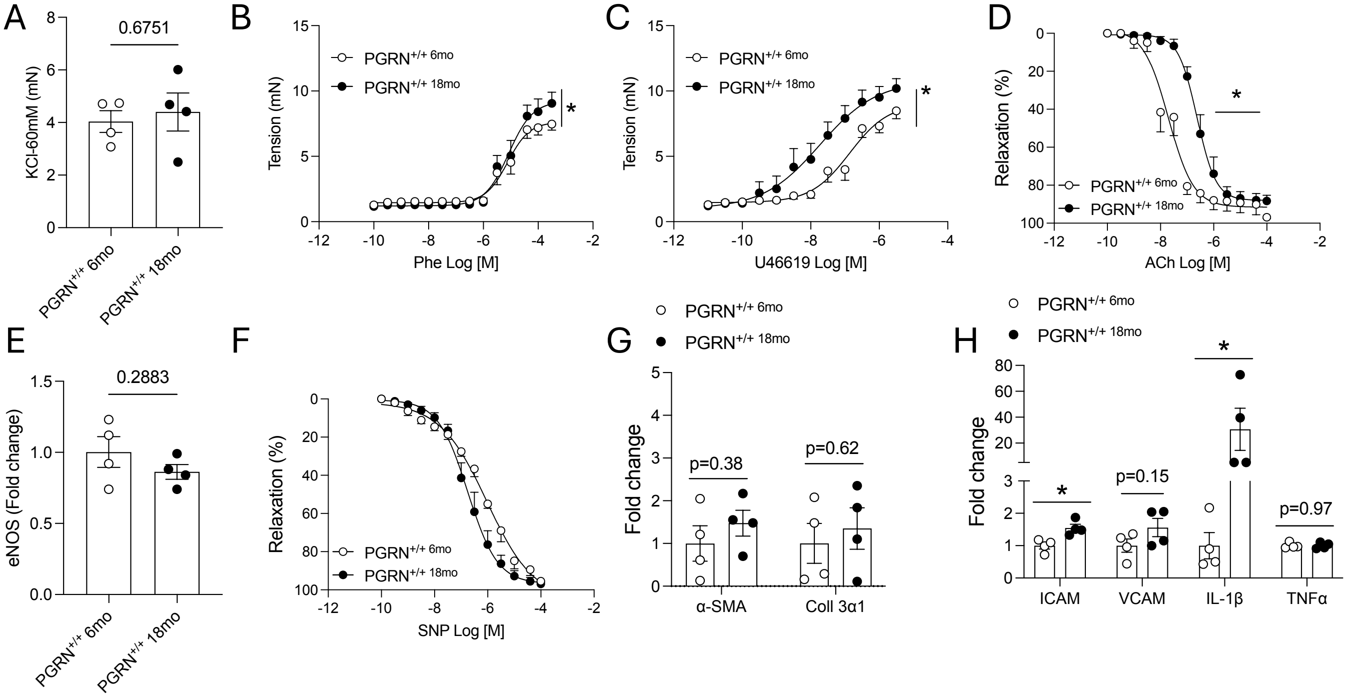


**Supplementary Figure 2. Aging impacts vascular function in mice.** (A) KCl-induced contractility. (B–C) Concentration–response curves (CRCs) to phenylephrine and U46619. (D) CRC to acetylcholine (ACh). (E) eNOS mRNA expression. (F) CRC to sodium nitroprusside (SNP). (G–H) Expression of structural and inflammatory genes. Experiments were conducted in mesenteric arteries from 6- and 18-month-old PGRN+/+ mice. Data are shown as mean ± SEM, N = 4. *P < 0.05 vs. PGRN+/+.

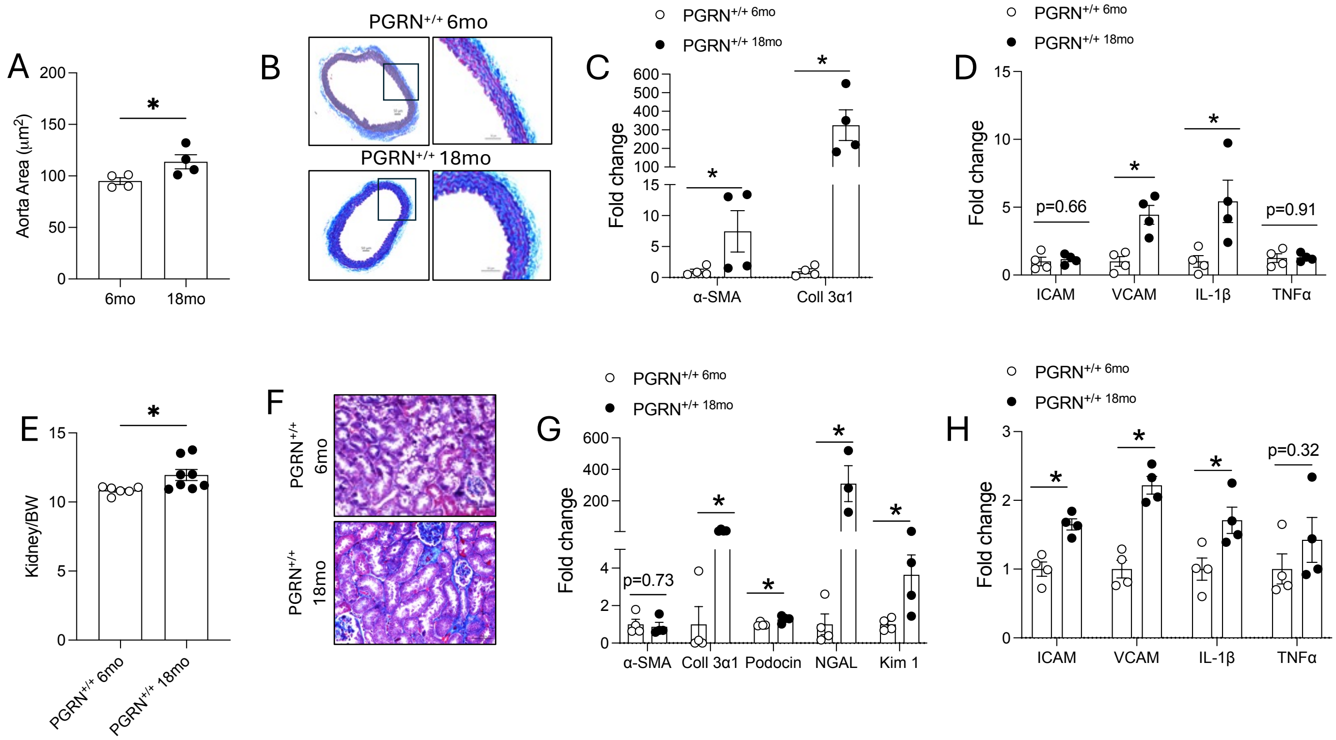


**Supplementary Figure 3. Aging exacerbates end-organ injury in mice. (**A) Cross-sectional area of the aorta. (B) Masson's Trichrome staining of aorta. (C–D) Expression of structural and inflammatory genes in aorta. (E) Kidney weight. (F) Masson's Trichrome staining of kidneys. (G–H) Expression of structural and inflammatory genes in kidneys. Experiments were performed in 6- and 18-month-old PGRN+/+ mice. Data are shown as mean ± SEM, N = 4-8. *P < 0.05 vs. PGRN+/+.

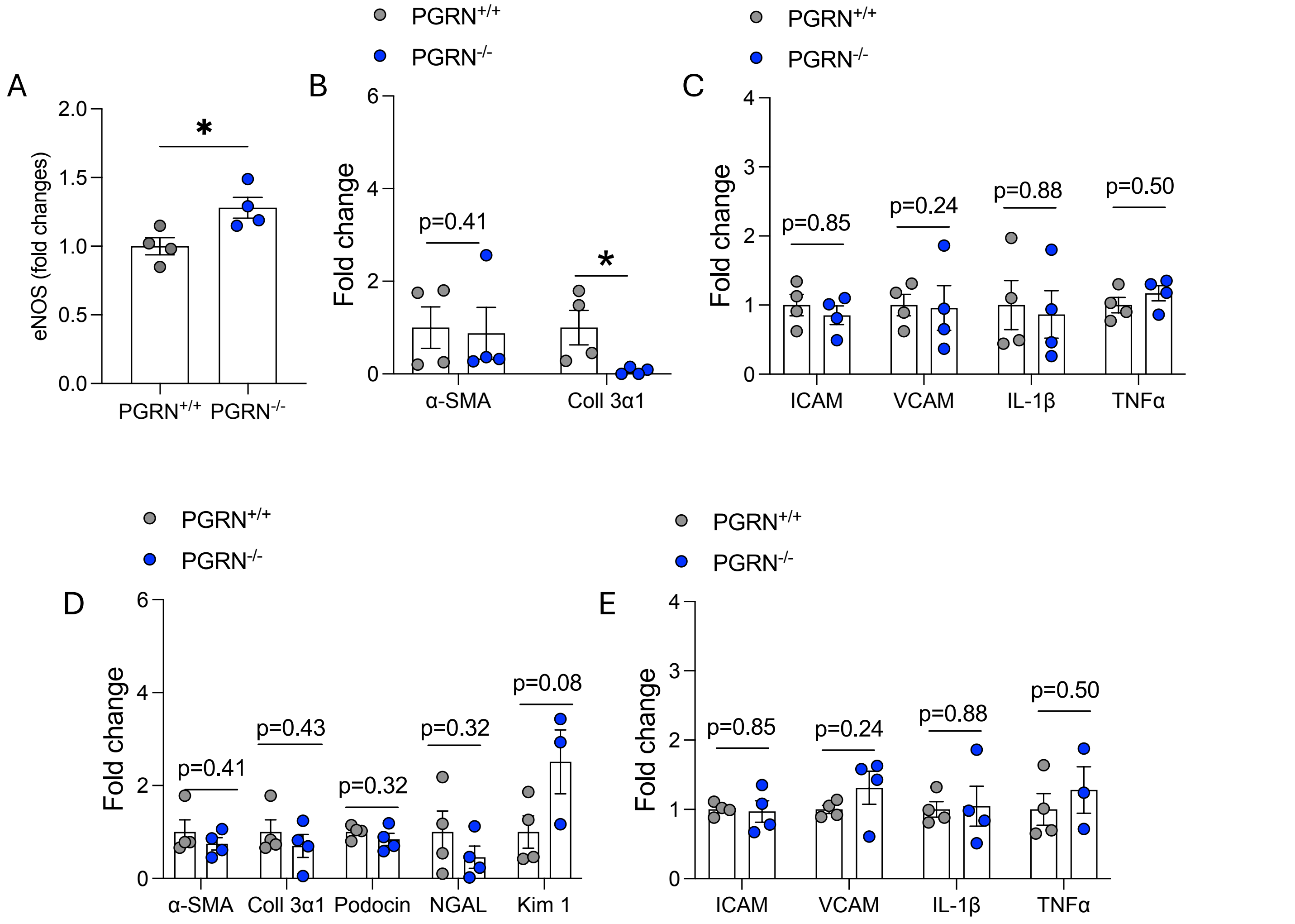


**Supplementary Figure 4. Progranulin (PGRN) deficiency does not further increase RNA markers of end-organ damage in aged mice.** (A) eNOS mRNA expression in mesenteric arteries. (B-C) Expression of structural and inflammatory genes in aorta. (D–E) Expression of structural and inflammatory genes in kidneys. All experiments were performed in samples from 18-month-old PGRN+/+ and PGRN-/- mice. Data are shown as mean ± SEM, N = 4. *P < 0.05 vs. PGRN+/+.
